# Mu oscillations and motor imagery performance: A reflection of success, not ability

**DOI:** 10.1101/2020.09.21.291492

**Authors:** Yvonne Y Chen, Kathryn Lambert, Christopher R Madan, Anthony Singhal

**Author notes:** Corresponding author, S101 Smith Research Building, One Baylor Plaza, Houston, TX, USA, 77030. both authors contributed equally (Yvonne Y Chen, Kathryn Lambert).

## Abstract

Motor imagery, or our ability to imagine movement without actually engaging in the action, has been an increasingly popular tool in rehabilitation settings. Understanding its neural underpinning is crucial for further development of new interventions. Using scalp electroencephalography (EEG), many studies have shown that mu oscillations (8-13 Hz), a variant of the alpha band recorded over the motor cortex electrodes, are involved in both the imagination and performance of movements; however, the exact relationship between mu oscillations and motor imagery is unclear. To further our understanding of the functional significance of mu oscillations and their role in both motor learning and motor performance, our study sought to investigate how suppression in mu oscillations varies during a motor imagery task according to both within subject imagery success and between subject imagery ability. We examined EEG activity while a large sample of participants performed an objective test of motor imagery ability (Test of Ability in Movement Imagery, TAMI). Results demonstrated that mu oscillatory activity significantly decreased during successful imagery trials as compared to unsuccessful ones. However, the extent of reduction in mu oscillations did not correlate with individual imagery ability. These results provide further support for the involvement of mu oscillations in motor behaviours and indicate that suppression in mu oscillations may serve as an important index for determining successful motor imagery performance within an individual. The processes that underlie this success are likely similar to those that underlie successful motor execution, given motor imagery’s proposed functional equivalence to motor imagery.

## 1. Introduction

Motor imagery refers to the mental rehearsal of movement from the first-person perspective in the absence of overt motor action (Jeannerod 1995; Madan & Singhal, 2012; Munzert et al., 2015; Munzert et al., 2009). Indications that this cognitive strategy can facilitate motor skill acquisition and enhance motor performance have made it a popular research area in recent years, with particular interest being directed to possible applications in both athletic and clinical settings. For example, the use of motor imagery based training as a rehabilitation tool for stroke patients suffering from hemiparesis has been researched extensively (for a review, see Page, 2010).

Motor imagery is thought to engage a similar neural network to motor execution. Neuroimaging research has shown that both motor imagery and execution engage a large fronto-parietal network and several subcortical regions, including the basal ganglia and the cerebellum (Hétu et al., 2013). These areas are involved in motor control and motor planning, both of which are necessary to the successful imagination and execution of movements. However, the exact pattern of neural activation during motor imagery varies based on both the nature of the imagined task and the nature of the imagination itself (Hétu et al., 2013). An imagined task can be as simple as a finger tap or as complex as a professional ballet routine, and while instruction can be provided on how the task should be imagined, the internal nature of motor imagery makes this difficult to enforce. It is likely that no two individuals imagine a movement exactly alike.

The process of motor imagery is often accompanied by visual imagery, the generation of a mental image without external stimuli (Kosslyn et al., 2010). Despite their regular co-occurrence, motor and visual imagery are considered two distinct processes, with both being forms of a more general ability known as ‘mental imagery’ (Kosslyn et al., 2010). Motor imagery is generally from a first-person perspective and involves dynamic images, whereas in visual imagery, the images are static and visualized from the third-person perspective (Madan & Singhal, 2012).

Both forms of imagery have been studied extensively using electroencephalographic methods, with this research focusing primarily on the alpha and mu rhythms. While these two rhythms oscillate within the same frequency range (8-14 Hz), they can be differentiated based on their distinct topographic and functional characteristics. The alpha rhythm is recorded over the centro-occipital region and dominates the brain at rest. It has been widely implicated in various cognitive processes, and is proposed to reflect an attention gating mechanism, wherein decreases in alpha activity reflects an increase in attentional demands (Jensen & Mazaheri, 2010). More importantly, decreased alpha activity, recorded from the parieto-occipital regions, has also been observed in many visual imagery tasks (Davidson & Schwartz, 1977; Barrett & Ehrlichman 1982, 1983; Golla et al., 1943; Marks & Isaac, 1995; Neuper et al., 2005). The decrease in alpha activity over the parieto-occipital regions may reflect active engagement of visual cortex during visual imagery (Salenius et al., 1995).

When tasks combine both visual and motor imagery, decreased activity is found not only in the alpha rhythm, but also in mu (Davidson & Schwartz, 1977). In contrast to alpha, the mu rhythm is recorded from the sensorimotor region of the human scalp (corresponding to paracentral areas) and consistently decreases during motor related processes (Neuper & Pfurtscheller, 2010; McFarland et al., 2000). This decrease typically occurs in the cortex that is contralateral to the performed movement and is localized to the sensorimotor region dedicated to the body part being moved (Pfurtscheller & Neuper, 1997; McFarland et al., 2000). It has been well documented that the same pattern of decreased mu activity occurs in motor imagery, which is not surprising given that motor execution and imagery share much of the same neural network (Davidson & Schwartz, 1977; Pfurtscheller & Neuper, 1997; McFarland et al., 2000; Formaggio et al., 2010). However, the decreases in mu that accompany motor execution are much greater than those accompanying motor imagery (McFarland et al., 2000).

While it is clear that decreased mu activity is related to motor imagery, the extent to which this decrease is related to imagery quality or capability remains unclear. ter Horst et al. (2013) explored the process by which mu might facilitate motor imagery through a mental hand rotation task. Participants were instructed to imagine rotating their hands both medially and laterally. While decreases in mu occurred in both the medial and lateral rotation tasks, the decrease was particularly intense in the medial condition. ter Horst et al. speculated that this difference was due to participant familiarity with medial hand rotation, which is more commonly performed in real life. Participants would therefore have had a stronger motor framework to draw upon during the imagery process. It was additionally proposed that the mu rhythm may act as a marker for motor imagery success, with greater decreases indicative of better imagery performance and more motor experience. Indeed, training protocols using brain computer interfaces (BCIs) have indicated that successful motor imagery is accompanied by greater decreases in the mu rhythm (Pichiorri et al., 2015; Ono et al., 2018).

While these findings suggest that greater decreases in mu should be observed individuals with higher motor imagery ability, given they are by definition more successful at imagery, no such correlation has been observed in recent research (Vasilyev et al., 2017; Toriyama et al., 2018). However, in an interesting result, Toriyama et al. (2018) found that differences in mu activity between motor execution and motor imagery were significantly less pronounced in vivid imagers. This suggests that the more vividly an individual imagines a movement, the closer it approximates his or her neural representation of motor execution. It is possible that comparisons between individuals is limited by overall variability in neural rhythms in the population (Blankertz et al., 2010). Further, a subset of individuals do not display decreases in mu activity during motor imagery, and levels of the mu rhythm at resting state vary significantly between individuals (Allison & Neuper, 2010; Blankertz et al., 2010).

A popular method for investigating imagery ability is through the administration of psychometric questionnaires that ask individuals to imagine the performance of particular movements and then rank the vividness of this imagery on a numeric scale (for a review, see McAvinue & Robertson, 2008). Research using vividness questionnaires suggests motor imagery ability can be improved with training and varies widely between individuals, including as a function of age, athletic skills, and overall health (Mulder et al., 2007; Eton et al., 1998; Ewan et al., 2010; Malouin et al., 2009; Malouin et al., 2008). However, as these questionnaires are self-report measures, their results may be confounded by several factors. Individuals may misestimate their ability due to over or under confidence, have differing interpretations of vividness levels, or feel pressured to report a certain score.

To avoid these possible biases, various objective measures have been explored as alternatives for measuring motor imagery ability. One such example is The Test of Ability in Movement Imagery, otherwise known as the TAMI (Madan & Singhal, 2013). In the TAMI, individuals are required to imagine a sequence of basic body movements and then select the body’s final positioning from five candidate images. Each question therefore has explicit correct and incorrect responses that are not subject to the biases of self-report questionnaires. Correlational investigations suggest that individuals engage in a first person perspective when performing the TAMI, with a greater emphasis placed on imagining how the movement looks to them as opposed to how it feels, the latter of which is known as kinaesthetic imagery (Madan & Singhal, 2013).

In addition to selecting an objective measure of motor imagery, we also chose an oscillatory quantifying measure that would be relatively selective for rhythmic activity and minimally influenced by non-repeating signals. A method for detecting oscillations, BOSC (Better OSCillation detection; Whitten, et al., 2011; Caplan, et al., 2001) is relatively conservative in its detection of rhythmic activity, with the results more specific to oscillations and less influenced by fleeting increases in power (Chen & Caplan, 2017). This method provides a measure of duration (Pepisode) that indicates the presence of oscillations at the chosen frequency during a specified time window. Using the measure of duration alongside a more conventional power (amplitude) measure of oscillations therefore enabled us to better evaluate the functional significance of mu oscillations in motor imagery.

In the current study, our main hypothesis was that the mu rhythm acts as an index of motor-imagery success within-individuals, wherein greater suppression indicates successful imagery performance. However, given significant variability of neural rhythms within the population, it is unlikely that this index of success translates to between-individual comparisons. We therefore hypothesized that differences in mu oscillations between successful and unsuccessful trials could explain individual motor imagery ability measured by TAMI. We recruited a large sample of EEG participants to explore the relationship between mu oscillations and their role in individual differences in motor imagery ability. In addition, visual imagery as reflected by decreased alpha activity is also anticipated to accompany motor imagery success.

## 2. Methods

### 2.1 Participants

A total of 80 introductory psychology students (58 female, 78 right-handed as measured by Edinburgh Handedness Inventory, Oldfield, 1971), aged 17 – 34 (M=19.43, SD=2.62) at the University of Alberta participated in the experiment for partial course credit. Data from 23 participants were excluded from analysis: 21 due to excessive EEG artefacts, and two due to handedness (score of < 40 on the Edinburgh Handedness Inventory, indicating ambidextrous or left-handedness). All participants had normal or corrected-to-normal vision. Written informed consent was obtained prior to the experiment in accordance with the University of Alberta’s ethical review board.

### 2.2 Procedure

The experiment was created and run using E-Prime presentation software version 2.0 (Psychology Software Tools). Participants completed a computerized version of the Test of Ability in Movement Imagery (TAMI) in an electrically shielded, sound attenuated chamber. The images and questions were presented in the center of the computer screen on a white background and a practice question preceded the start of the experiment to familiarize participants with the format of the TAMI. Participants were able to proceed with the test upon answering the question correctly.

Participants completed 10 questions of the TAMI, with each question made up of four separate movements. Each question begins with the identical instruction, asking the participant to imagine the following starting position: “Stand up straight with your feet together and your hands at your sides”. This instruction was presented alongside an image showing this body position (Starting position, Step 0). The subsequent four steps differed with every question, but each required the individual to imagine the movement of a specific body part. Each step was released after a 6000 ms delay, giving participants time to imagine the movement. Once the last step was presented, a response screen would appear and present a set of five images showing the possible final body position, along with the choices of “none of the above” and “unclear”. The images remained onscreen until the participants made a response by pressing the key corresponding to the image of their choice. Refer to Figure 1 for a visual of the procedure.

**Figure 1.**
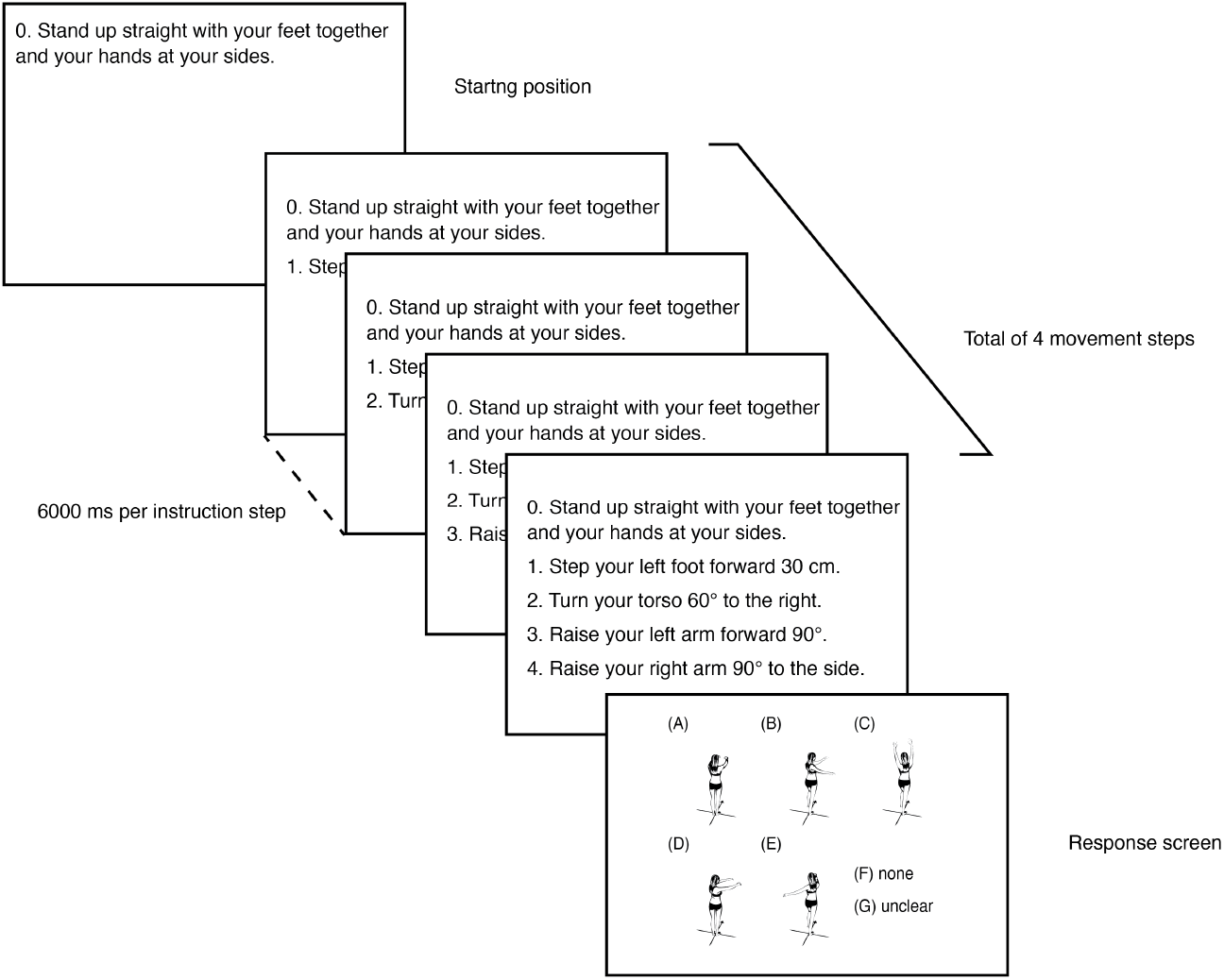
The experimental procedure. Each box illustrated the computer screen at a particular stage in the task (texts and images have been enlarged relative to the screen size to improve clarity of the figure. Each question starts with an initial standing position followed by five instruction steps. Participants respond by pressing a response button corresponding to their imagined final body positioning at the “response screen”. There are a total of 10 imagery questions.

### 2.3 EEG recording and analyses

EEG was recorded using a high-density 256-channel Geodesic Sensor Net (Electrical Geodesics Inc., Eugene, OR), amplified at a gain of 1000 and sampled at 250 Hz. Impedance was kept below 50kΩ and recording was initially referenced to the vertex electrode (Cz). Data was analyzed by custom MATLAB scripts in conjunction with open-source EEGLAB toolbox (Delorme & Makeig, 2004). The signal was bandpass filtered from 0.1 to 50 Hz and average referenced. Artifacts were corrected via independent component analysis, implemented in EEGLAB. The selection of components was based on visual inspection of the spatial topographies, time courses, and power spectral characteristics of all components (Jung et al., 2000). The components accounting for stereotyped artifacts including eye blinks, eye movements, and muscle movements were removed from the data. Event latencies were corrected with a time lag correction due to a known hardware calibration problem identified by EGI.

### 2.4 Spectra power and duration (Pepisode) analysis

The continuous (unepoched) preprocessed signal was then analyzed for oscillatory activity using the power and duration measure (Better OSCillation detection method, Caplan et al., 2001; Whitten et al., 2010). First, the preprocessed EEG-signal was convolved with a Morlet wavelet transform with a width of six cycles and sampled at 24 frequencies logarithmically over 1 - 45 Hz range. Power measures were estimated from the square of instantaneous amplitude of the complex convolution results, and then log-transformed and normalized by dividing the mean log-power from the entire recording session at each frequency band and at each electrode. BOSC method classifies oscillatory events when a segment of signal power at a given frequency exceeds the power threshold for a minimum duration of time (duration threshold). Thus, we set a power and duration threshold to ensure we captured true rhythmic activity. To this end, the power spectrum threshold was calculated to exclude 95% of the signal’s background, or coloured noise, from analysis. We set a duration threshold of three cycles to ensure that fleeting increases in power were not considered oscillatory. The result obtained from this method is a duration measure (Pepisode), which represents for a given frequency, the duration (proportion) of time at a given interval detected by BOSC. The measure ranges from 0 to 1, where Peipisode = 0.5 means that a given frequency was detected in half of the analysis.

We selected four electrodes for analysis, each of which corresponded to a brain region of interest. The C3, C4 and Cz are the principal electrodes for the motor area. All found over the brain’s central region, C3 and C4 are located on opposite sides of the scalp, left and right respectively. Several studies have reported mu oscillations from C3 and C4 (Formaggio et al., 2010; Pfurtscheller & Neuper, 1997; Pfurscheller et al., 2006; Pfurtscheller et al., 2005; Nam et al., 2011; McFarland et al., 2000). Analysis of these electrodes enabled us to make comparisons with findings from similar studies. By also selecting Oz for analysis, a primary electrode for the alpha rhythm, we were able to determine how activity in the mu rhythm differed from other rhythms in the same frequency band.

Our analysis was concentrated on comparison of activity between unsuccessful and successful trials within the same participant. Given that individuals (N=7) who scored a 10 out of 10 on the TAMI questions had no unsuccessful trials for comparison, their data was excluded from this analysis, reducing the total number of participants to 53. For this approach, mu and alpha frequency bands were defined as 8-14 Hz. Frequencies within each band were collapsed by averaging together the Pepisode values obtained within its previously specified bandwidth. The P_episodes_ were then averaged together across participants, categorized according to condition (successful vs. unsuccessful trials) and compared using a paired-sample, two-tailed t-test.

## 3 Results

The primary goal of this study was to investigate if mu oscillations index motor imagery success and ability as measured by an objective imagery assessment tool, the TAMI. We first examined imagery success by contrasting successful and unsuccessful motor imagery trials within a given participant, predicting that less mu oscillations would accompany the successful trials. Next, we investigated individual imagery ability by correlating mu oscillations with performance on the TAMI score across participants. If mu suppression is a reflection of an individual’s ability to perform motor imagery, we would expect individuals who score higher on the TAMI to have a greater magnitude of difference in mu oscillatory activity between successful and unsuccessful trials, indicating that their successful performance was accompanied by more suppression in the mu rhythm.

### 3.1 Behavioral Results

We first analyzed participants’ performance on the TAMI. The mean score was 7.4 (SD= 1.75) out of 10. This score is comparable to the original TAMI score reported in Madan and Singhal (2013). Given our sample was comprised primarily of young, healthy individuals, we also considered the adjusted weighting scoring developed by Madan and Singhal (2014). This method helps to reduce the ceiling effect that often accompanies high performance on tasks. The mean weighted score of our participants was 15.6 (SD = 5.19) out of 24. Figure 2 illustrated the distribution of participants’ TAMI scores, unweighted (A) and weighed (B).

**Figure 2.**
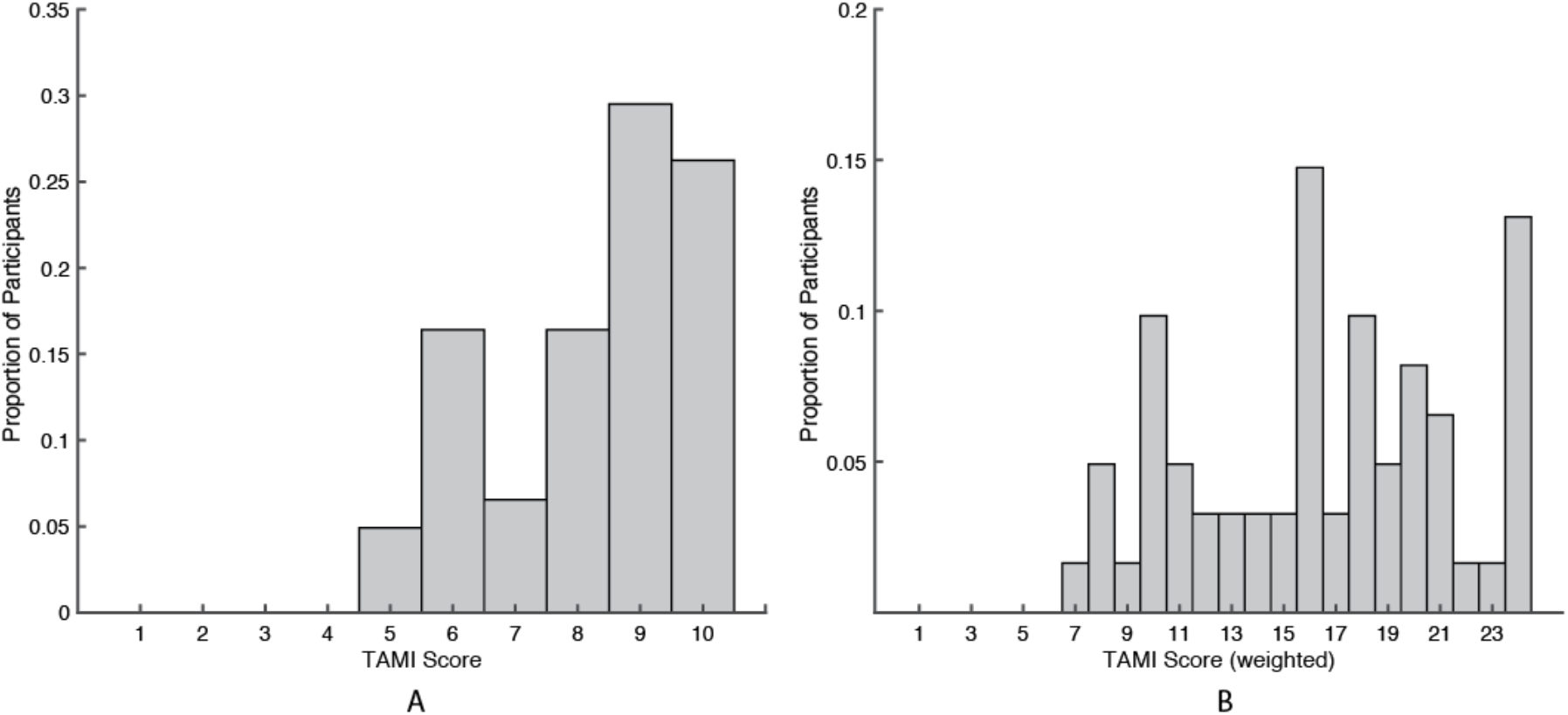
Distribution of participants’ scores on the Test of Ability in Movement Imagery (TAMI), (A) unweighted and (B) weighted TAMI score.

### 3.2 Oscillation Results

To test the hypothesis that suppression in mu activity, as demonstrated by fewer oscillations detected, facilitates successful motor imagery, we compared oscillatory activity between successful and unsuccessful trials within each participant. Trials were scored as “successful” only when participants chose a single correct response to the question. Mu oscillations were quantified using both Power and Pepisodes, a proportional duration measure, at the central electrodes (C3, C4 and Cz). Our time window of analysis was each TAMI question in its entirety, a twenty four second “imagery” period that began with the onset of the first movement step and concluded with the offset of the fourth movement step (6000 ms per step). The period during which participants selected their response was not included in this analysis as our focus was on the process, not the end product, of motor imagery. The measures of oscillations (both Pepisode and power) are shown in Figure 3. Apart from central motor electrodes, we also examined alpha activity over the occipital region (electrodes Oz) due to the rhythm’s possible role in motor imagery.

**Figure 3.**
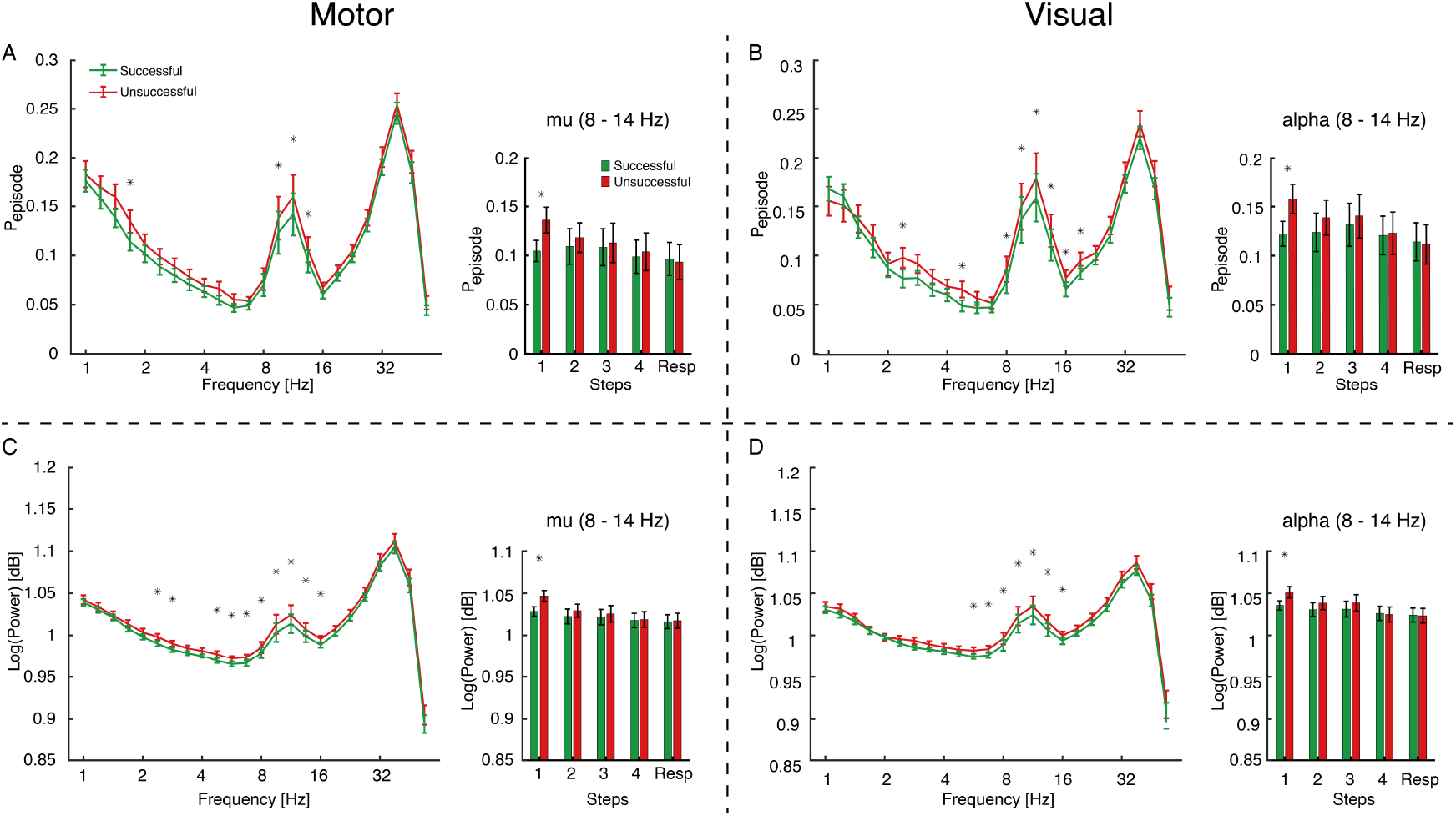
Oscillatory activity and motor-imagery success. Group average proportion of oscillatory activity (Pepisode) and group average wavelet power (log-transformed) for entire TAMI question are plotted as a function of frequency between successful (green) and unsuccessful (red) trials. Mu and alpha oscillations are plotted as a function of movement steps (steps 1-4 and response screen) between successful (green) and unsuccessful (red) trials. Error bars represent 95% confidence intervals and * denotes a pairwise t test significant (p < 0.05) difference between successful and unsuccessful imagery trials.

A repeated-measure ANOVA comparing mu oscillations of imagery success (successful/unsuccessful) * electrode locations (C3/C4/Cz/Oz) revealed a main effect of imagery success as measured by Pepisode. F(1,52) = 18.91, p <0.001, due to less presence of mu oscillations during successful trials. A main effect of electrode locations was also significant, F(3,156) = 4.087, p <0.05. A post-hoc pairwise comparison (Bonferroni corrected) revealed no significant effect between central motor electrodes (C3, C4, and Cz), but there were significant differences between the motor electrodes and Oz, the visual electrode (C3 vs. Oz: t = −2.692, p(bonf) <0.05; C4 vs. Oz: t = −3.112, p(bonf)<0.05; Cz vs. Oz: t = −2.691, p(bonf)<0.05). We therefore grouped all central motor electrodes together by averaging across the three electrodes for the rest of our analysis. Another repeated-measure ANOVA comparing imagery success (successful/unsuccessful) * electrode locations (motor/visual), revealed a main effect of imagery success as measured by Pepisode F(1,52) = 15.35, p<0.01; and a main effect of electrode location F(1,52) = 11.84, p=0.01. The two main effects survived the post hoc pairwise bonferroni correction. The interaction was not significant (p>0.1). Similar results were also found using the same repeated-measure ANOVA examining imagery success measured by wavelet power and electrode locations. There was a main effect of imagery success by power F(1,52) = 12.793, p<0.01 and a main effect of location F(1, 52) = 8.431, p < 0.05. These two main effects survived the post hoc pairwise Bonferroni correction. The interaction of imagery success * electrode locations was not significant (p>0.1).

Together, these results suggest that less mu oscillations were present during successful motor imagery trials, indicating mu suppression reflects successful motor imagery. While we did not specifically focus on oscillations outside the mu frequency band, the results are not surprising. For example, Harmony (2013) has associated the delta band with attention to internal processing, a form of mental processing that is presumably required to perform mental imagery. Additionally, both the theta and beta rhythms have been implicated in motor processes, with the former linked to sensorimotor integration and the latter to motor imagery itself (Cruikshank et al., 2011; Formaggio et al., 2010; McFarland et al., 2000).

In addition, we were also interested in how mu oscillations change over time by comparing mu oscillations between successful and unsuccessful trials at each of the four movement steps and at the response page at motor and visual electrodes (Figure 3). A repeated three ANOVA comparing imagery success measured by Pepisode (successful/unsuccessful)*Steps (1/2/3/4/Resp)*electrode locations (motor/visual) revealed a main effect of imagery success, F(1,52) = 7.258, p<0.01, and a main effect of Steps, F(4,208) = 3.609, p <0.01, and a main effect of location, F(1,52) = 12.845, p<0.001. There was also a significant imagery success * steps interaction F(4,208) = 5.023, p<0.001 and there was no significant interaction of steps*location, imagery success*location, and imagery success*steps*location. A post hoc analysis using pairwise Bonferroni correction revealed that only the main effects of imagery success and locations were significant. In addition, only mu oscillations measured at Step 1 significantly differed between successful and unsuccessful trials at the motor (t = −4.306, p(bonf)<0.01) and visual (t = −4.887, p(bonf)<0.001) electrodes. The same repeated ANOVA using a power as opposed to duration measure revealed similar main effects and interactions. These results suggest that initial mu suppression at the onset of the motor imagery process is crucial to imagery success and that mu oscillation suppression is less sustained than perhaps previously thought.

We next tested our second hypothesis: that suppression of mu oscillations may index individual motor imagery ability. We quantified the mu oscillation suppression using the difference measure by calculating mu oscillation Pepisode difference between successful and unsuccessful trials within individuals. We chose to examine this difference measure of mu oscillations, because the difference measure infers the relationship between one’s ability to suppress mu oscillations and motor imagery ability, indicating that a high-motor-imagery-ability individual may possess a greater ability to suppress mu oscillations. To test this possible behavioral-relevance of mu oscillations, we first correlated across participants the difference in mu oscillation suppression (successful - unsuccessful) with TAMI scores (both weighted and unweighted). We found no significant correlation between mu oscillation suppression measured by Pepisode and TAMI scores [unweighted: motor: r(51) = −0.003, p >0.1; visual r(51) = 0.128, p>0.1; and weighted: motor: r(51) = 0.136, p >0.1; visual r(51) = 0.247, p>0.1]. Since we found suppression of mu oscillations to be strongest at the initial movement step, we also examined the relationship between mu oscillations from step 1 with TAMI scores. We correlated the mu oscillation difference measure (successful-unsuccessful) that occurred during the step 1 time window alone with TAMI scores (both weighted and unweighted). There remained no significant correlation [unweighted: motor: r(51) =- 0.015, p >0.1; visual r(51) = −0.012, p>0.1; and weighted: motor: r(51) = 0.115, p >0.1; visual r(51) = 0.141, p>0.1]. In addition, we also performed the same correlations using the wavelet power measure, and again found no significant correlations.

These null correlations suggest that the individual differences in mu oscillations, measured by difference between successful and unsuccessful trials, do not reflect individual motor imagery ability as measured by the TAMI. Apart from the central motor electrodes, when we examined alpha activity over the occipital region (electrodes Oz) using the same analysis approach, we found results similar to those of the mu oscillations recorded from motor electrodes. This suggests that alpha oscillations are also not a reflection individual motor imagery ability.

## 4 Discussion

The aim of the current study was to examine how oscillatory activity in the mu rhythm varies according to motor imagery success and individual motor imagery ability. We found that successful responses on an objective motor imagery task, the TAMI, were accompanied by greater decreases in mu oscillatory activity as compared to unsuccessful responses. However, this decrease in activity with successful performance did not correspond to differences in individual motor imagery ability. Individuals who received higher scores on the task, and were therefore considered to be of high imagery ability, did not have significantly different levels of mu activity when compared to those who performed poorly.

We were able to make confident comparisons between successful and unsuccessful performance due to the objective nature of our measure, with participants required to make explicitly correct or incorrect responses during the task. A key result therefore was the difference found between successful and unsuccessful responses, with less mu oscillations present during imagery that would later generate a successful response. This difference in activity between successful and unsuccessful responses was anticipated based on the findings of previous research (ter Horst et al.., 2013; Pichiorri et al., 2015; Ono et al., 2018). Greater decreases in mu activity have been found to accompany motor imagery of more familiar movements, movements which are likely easier for individuals to picture and thus more successfully imagined (ter Horst et la., 2013). Further, Brain Computer Interface (BCI) research has demonstrated that corrective feedback during imagery, designed to increase the accuracy of the imagery process, leads to reduced mu activity (Pichiorri et al., 2015; Ono et al., 2018). This supports the theory that mu oscillations reflect motor imagery success, a finding which may be especially useful in terms of motor imagery training protocols. The mu rhythm could possibly be used as a neurophysiological marker to indicate if training has helped an individual become more successful at motor imagery.

However, it is likely that our results provide only a limited picture of motor imagery success. Guillot et al. (2010) proposed motor imagery ability should be considered in terms of three separate dimensions: vividness, controllability and exactness. Vividness refers to the sensory richness of the self-generated image and exactness the image’s accuracy, while controllability reflects the ease of the image’s manipulation (Guillot et al., 2010). While these dimensions overlap, and are thus difficult to fully disentangle, different tasks emphasize different dimensions. For example, as the TAMI emphasizes accurate responses, it places more precedence on exactness than self-report vividness questionnaires. However, this does not mean that the vividness dimension goes ignored by the TAMI. Indeed, performance on the TAMI correlates with two different measures of self-report imagery vividness, the internal visual imagery subscale of the VMIQ and the VVIQ (Madan & Singhal, 2013). Further, the requirement that individuals manipulate an image during the TAMI means that some skill in controllability is also required. While the TAMI provides a comprehensive, objective measure of motor imagery, our results may not necessarily be reproduced by a measure that prioritized different imagery dimensions.

It is important to note that the question of which specific dimensions of motor imagery take precedence will varies within a task itself. Vividness and exactness are both key at the onset of the task, when the image is first being generated, as opposed to controllability, which is central to the image’s subsequent maintenance and transformation (Guillot et al., 2010). Interestingly, the significant difference in mu oscillatory activity found between successful and unsuccessful responses occurred only at the first of the four motor imagery steps of the TAMI. The initial step of the TAMI occurs immediately after the presentation of the starting body position, and is the first moment wherein the participants are required to manipulate an image in their minds. This result would suggest it is mu activity at the onset of the motor imagery that is essential to successful performance as opposed to its activity throughout the entire imagery process. Perhaps the mu rhythm reflects the generation of the initial image, with greater decreases in oscillatory activity the reflection of a more accurate, vivid image, which in turn leads to a greater chance of later motor imagery success.

If successful responses within individuals result in decreased mu activity, it would make intuitive sense that good imagers, who by definition have more successful responses on the TAMI, would have less mu oscillatory activity overall than poor imagers. However, this was not the case, and our previous results were not echoed in the between-subjects comparison. While both good and poor imagers exhibited decreases in mu oscillatory activity during successful trials, the good imagers demonstrated no less overall mu activity than their poorer scoring counterparts. Our result is in line with recent research, wherein individuals with higher scores on the vividness self-report questionnaires, an indication of higher imagery ability, did not exhibit greater decreases in mu activity as compared to those with low scores (Vasilyev et al., 2017; Toriyama et al., 2018). This lack of difference in oscillatory activity as a function of overall motor imagery ability may be a result of the rhythm’s quantitative instability and variability across the population, with mu activity levels at baseline differing between individuals (Allison & Neuper, 2010; Blankertz et al., 2010). When taken into consideration with prior findings, our results provide further indication that the mu rhythm alone should not be used to index overall motor imagery ability, but rather as a possible measurement of success within an individual.

Beyond mu oscillations, we also found less alpha activity as recorded over the occipital region during successful motor imagery when compared to unsuccessful motor imagery. Alpha oscillations have been linked to a variety of cognitive tasks, including attention (Hanslmayr et al., 2011, Foxe and Snyder, 2011; Jensen and Mazaheri, 2010), working memory (Jensen et al., 2002; Klimesch et al., 2006), and visual imagery. Alpha activity has traditionally been thought to reflect cortical activation (Klimesch, 1999; Klimesch, 2012). More recently, the alpha rhythm was thought to reflect inhibition, with decreased alpha activity reflecting a release from inhibition that “sharpens” an individual’s attentional view and enables them to better process on task demands(Cooper et al., 2003; Klimesch, et al., 2007; Jensen & Mazaheri, 2010). Moreover, other research has also suggested alpha activity is sensitive to attentional demands. It is possible that the differences in alpha activity we observed between successful and unsuccessful responses is a reflection of attentional state, where more alpha activity inhibited the motor imagery process from fully taking place. In addition, alpha oscillations have also been suggested to reflect working memory function, wherein effortful attention is required to hold information in the mind. Indeed, activity in the alpha rhythm has been shown to decrease as working memory load increases (Gevins et al., 1997). We observed decreased alpha activity in the successful trials as compared to the unsuccessful trials. It is possible that participants were more successful in holding information in their mind when they generated the correct response.

Lastly, it is also important to consider the function of alpha oscillations in visual imagery tasks. There are two main oppositional findings of alpha oscillations and visual imagery. First, increases in alpha activity are associated with the imagery process. Many prior studies have suggested that alpha oscillations support the maintenance of internal representations, and are thus essential to the successful imagination of a visual object or a sequence of motor movements (see Klimesch et al., 2007, for a review). When Bartsch et al. (2016) asked participants to imagine visual objects based on the verbal instructions given, they found an association between increased alpha power and word-prompted mental imagery. However, decreases in alpha activity are also associated with the imagery process, stemming from the idea that visual perception and visual imagery may rely on similar neural substrates as indexed by alpha suppression (Lee et al 2012). For example, during a mental rotation task, Michel et al. (1994) found less alpha activity during the process of mental visual imagery. Although TAMI is a motor imagery task, it still heavily relies on visual imagery. It is likely that our participants may engage in visual imagery to aid their motor imagery process, and alpha oscillations measured over the occipital regions reflect the visual imagery process. While we may not be able to resolve the debate on alpha activity and motor imagery, we added additional evidence that reduced alpha activity is associated with objective success in motor imagery.

## 5 Conclusion

The present study provides evidence that mu oscillations and imagery success are related. Our finding of decreased mu activity during successful performance is in line with previous research linking the rhythm to imagery performance. However, no significant differences in the rate of occurrence of mu oscillatory activity were found between individuals of high imagery ability and those of lower imagery ability. This suggests that mu oscillations may index successful imagery performance within an individual but cannot be used to make comparisons across a population. Results in the occipital region suggest that visual imagery, as reflected by the alpha rhythm, may facilitate the function of motor imagery in TAMI. At present, we are unable to properly disentangle these two different forms of imagery (motor and visual). Nevertheless, our result confirmed the prior hypothesis of mu oscillations as an indicator of individual motor imagery success.

## Acknowledgments

The authors would like to thank Vivian Chan and Tristan Pasek for their help in collecting EEG data, and Drs. Ada Leung and Eleonora Bartoli for helpful comments on the manuscript. This research was partly funded by the Natural Sciences and Engineering Research Council of Canada, the Canada Foundation for Innovation, and the University of Alberta Neuroscience and Mental Health Institute.

